# Calcium oscillations in wounded fibroblast monolayers are spatially regulated through substrate mechanics

**DOI:** 10.1101/116426

**Authors:** Josephine Lembong, Benedikt Sabass, Howard A. Stone

## Abstract

The maintenance of tissue integrity is essential for the life of multicellular organisms. Healing of a skin wound is a paradigm for how various cell types localize and repair tissue perturbations in an orchestrated fashion. To investigate biophysical mechanisms associated with wound localization, we focus on a model system consisting of a fibroblast monolayer on an elastic substrate. We find that the creation of an edge in the monolayer causes cytosolic calcium oscillations throughout the monolayer. The oscillation frequency increases with cell density, which shows that wound-induced calcium oscillations occur collectively. Inhibition of myosin II reduces the number of oscillating cells, demonstrating a coupling between actomyosin activity and calcium response. The spatial distribution of oscillating cells depends on the stiffness of the substrate. For soft substrates with a Young’s modulus *E* ~ 360 Pa, oscillations occur on average within 0.2 mm distance from the wound edge. Increasing substrate stiffness leads to an average localization of oscillations away from the edge (up to ~0.6 mm). In addition, we use traction force microscopy to determine stresses between cells and substrate. We find that an increase of substrate rigidity leads to a higher traction magnitude. For *E* < ~2 kPa, the traction magnitude is strongly concentrated at the monolayer edge, while for *E* > ~8 kPa, traction magnitude is on average almost uniform beneath the monolayer. Thus, the spatial occurrence of calcium oscillations correlates with the cell-substrate traction. Overall, the experiments with fibroblasts demonstrate a collective, chemomechanical localization mechanism at the edge of a wound with a potential physiological role.

## INTRODUCTION

Wound healing in mammalian organisms is a complex process that requires coordination of various tissues and involves highly regulated activities, including cell proliferation and migration (1–3). In a dense group, cells exert forces on each other via cadherin-based adhesions (4). Simultaneously, they use integrin-based adhesions to exert forces on their surrounding extracellular matrix (ECM) (5–7). Together, ECM adhesions and cell-cell adhesions serve to maintain the integrity of cellular tissue while allowing both migration and adhesion (4, 8–12). Forces are produced mostly by the acto-myosin machinery, which is regulated by a plethora of signaling pathways, e.g., the prominent Rac and Rho pathway (13–16), but also by intracellular calcium (17, 18). Investigation of the response of such systems to mechanical perturbations is both a topical and fundamental research subject.

Typically, studies of collective cell mechanics in wound healing model systems employ epithelial cells (16, 19–23). Epithelial cells undergo multiple biochemical changes during wound healing that enable the acquisition of mesenchymal cell phenotypes, which includes enhanced migratory capacity, invasiveness, increased resistance to apoptosis, and amplified production of extracellular matrix components (24). During this epithelial-mesenchymal transition (EMT), fibroblasts are generated. Along with resident fibroblasts and other mesenchymal cells, EMT-derived fibroblasts play a key role during reconstruction of tissues following a wound or inflammatory injury (1, 24). Therefore, studying the collective response of fibroblast-like cells in a wounded model tissue promises to reveal novel biophysics with possible physiological implications. In the present study, we employ a fibroblast cell line.

Intracellular calcium is a regulator of cellular mechanics (25, 26). In motile fibroblasts, calcium acts through motor proteins such as myosin II, which are co-localized with calmodulin in the contracting tails during locomotion (27). Calcium-calmodulin can activate myosin light chain kinase (MLCK), which in turn activates myosin II, generating contractile force and hydrostatic pressure that can propel the cell forward. Furthermore, calcium acts on various actin-binding proteins, e.g. fodrin (28) and MARCKS (29), which exhibit decreased cross-linking activity in the presence of high calcium. Fibroblasts transfected with gelsolin, a calcium-regulated protein that severs actin, exhibit increased rates of motility (30). Additionally, changes in Ca^2+^ affects the cell adhesion state via surface integrins (31). In mouse fibroblasts, integrin engagement also results in generation of second messenger inositol triphosphate (IP3) and subsequently release of Ca^2+^ from internal Ca^2+^ stores (32). Through various signaling pathways involving Ca^2+^ such as the MAP kinase cascade, including those mediated by adhesion receptors of the integrin family and those mediated by serum (33, 34), regulation of cell mechanics, including motility and migration, can be achieved. Directionality of cell motion has also been linked to local Ca^2+^ pulses near the leading edge, which are involved in the activation of MLCK and modulation of focal adhesions (35–37).

Several studies have investigated the intracellular Ca^2+^ dynamics in the context of migrating cell ensembles. In various cell types cultured on glass substrates, levels of intracellular Ca^2+^ are observed to correlate with migration speed (table S1 in the supplementary material) (38–42), which are attributed to local modulation of the cytoskeleton at the integrin-mediated adhesion sites. However, the rigidity of the extracellular environment is clearly an important determinant of cellular behavior (43–47). For example, NIH 3T3 fibroblast cells are known to migrate from soft to rigid substrates (48). In addition, intracellular calcium oscillations in fibroblasts are known to be affected by the substrate mechanical properties (49, 50). A better understanding of the link between calcium dynamics and the cells’ mechanical environment can help decipher cellular processes underlying cell ensembles whose dynamics is constantly dictated by cell-cell and cell-substrate interactions.

In this study, we explore the phenomenology of the coupling between calcium oscillations and cellular forces after wounding a fibroblast monolayer. The cells are cultured on a glass substrate or on a polyacrylamide (PA) gel with Young’s moduli *E* between 360 and 25000 Pa, which mimics the order of magnitude of the elasticity of various epithelial and fibroblast tissue environments (51, 52). We observe intracellular calcium oscillations upon creation of an edge in the fibroblast monolayer. The oscillations occur on a timescale of minutes and the frequency increases with cell density. A rigidity-dependent localization of calcium oscillations in the fibroblast monolayer correlates spatially with cell-substrate traction. Increase of substrate rigidity leads to more oscillating cells and a higher average traction. The observed calcium dynamics of the fibroblast monolayers provide insight into a coupled chemical-mechanical mechanism for wound localization that may also occur during the initial stages of wound closure.

## METHODS

### Cell culture and substrate preparation

NIH 3T3 mouse fibroblast cells (ATCC, Rockville, MD, USA) were cultured as previously described (49). For cells grown on polyacrylamide (PA) gel (Bio-Rad, Hercules, CA, USA), activated glass-bottom petri dishes were first prepared to ensure binding of the PA gel to the glass surface. The treatment started with soaking glass-bottom petri dishes in 1% 3-aminopropyltrimethyoxysilane in deionized (DI) water for 10 min, followed by 10 min in DI water, 10 min in 0.5% glutaraldehyde in phosphate-buffered saline (PBS; ATCC) and finally 30 min in DI water. PA gel solutions (with 1 vol % fluorescent beads (*d* = 200 nm), 0.05 vol % ammonium persulfate, and 0.15 vol % tetramethylethylenediamine (TEMED)) in water were prepared using compositions listed in figure S1 and table S2 in the supplementary material. A small volume (approx. 50 μ1) of the PA solution mixtures of different ratios was placed on the activated glass-bottom petri dishes, then covered with hydrophobic coverslips (*d* = 25 mm). This volume resulted in PA gels with an area of ~500 mm^2^ and ~100 μm thickness. After the PA gel polymerized (approx. 20 min at room temperature), the hydrophobic coverslip was peeled carefully. To allow cell attachment to the PA substrate, fibronectin (Sigma-Aldrich, St Louis, MO, USA) was chemically cross-linked to the surface of the cured PA gel using the photoactivated cross-linker sulfo-SANPAH (Fisher Scientific, Pittsburgh, PA, USA).

### Creation of model wounds

Fibroblast cells were grown on glass bottom petri dishes (Ted Pella, Redding, CA, USA) or polyacrylamide (PA) gel surrounding a PDMS strip, with dimensions 25 mm × 2 mm × 2 mm (figure 1). Fibroblast cells were left to adhere on the substrates for 24 hours. Prior to imaging, Fluo-4 calcium dye (Invitrogen, Carlsbad, CA, USA) was administered in growth medium and left to incubate with the cell culture for 1 hour. The model wound was created by removing the PDMS strip (19), which exposed the cells to a free environment at the open edge and therefore created a change in the surrounding mechanical stresses. Upon PDMS removal, the cytosolic calcium level was measured around the area of the edge indicated by the red rectangle in figure 1(a).

**Figure 1.**
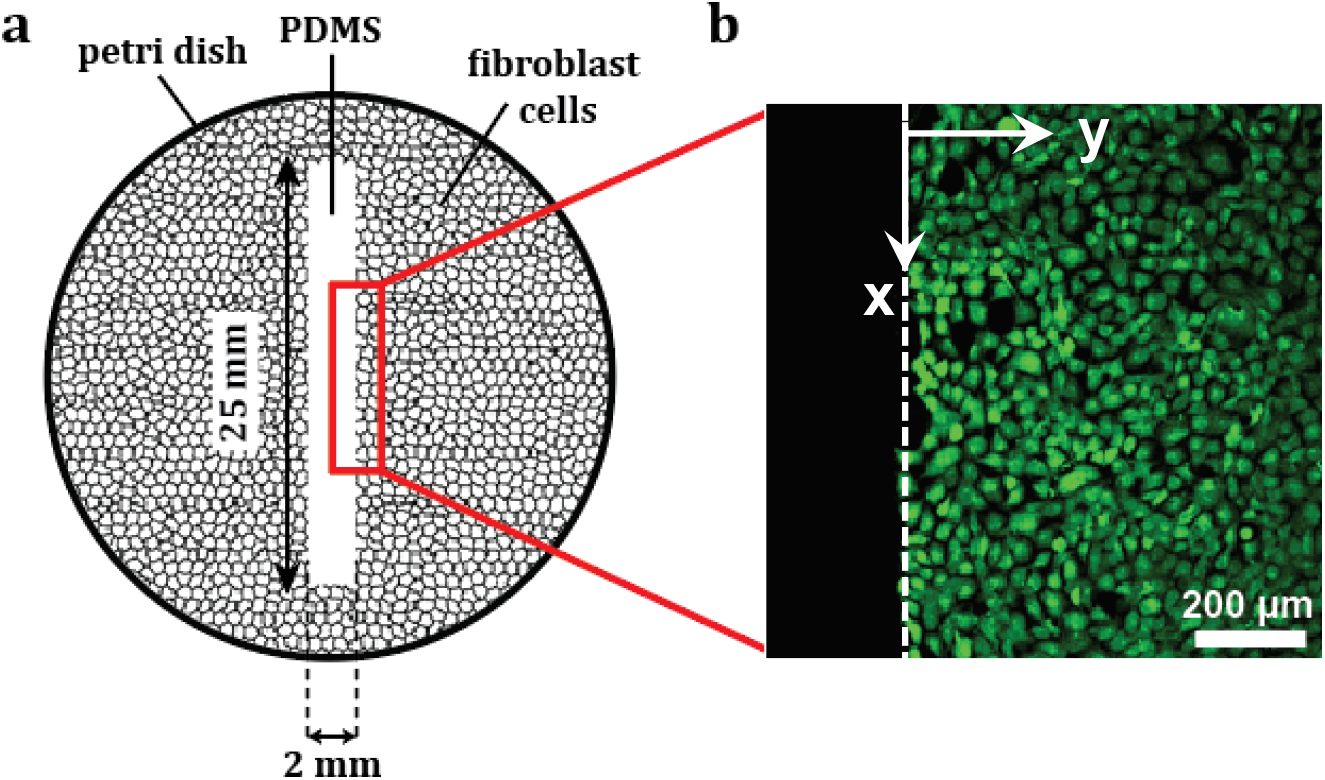
Schematic of the experimental “wound” setup. (a) Cells were cultured on a fibronectin-coated glass-bottom petri dish or PA gel around a PDMS strip (cells not drawn to scale). Imaging of the cytosolic calcium level started immediately following PDMS removal. (b) Fluorescent image of fibroblast cells near the free edge after wound creation.

### Fluorescence microscopy for Ca^2+^ imaging

Fluorescence was detected using a confocal microscope (Leica SP5; Leica Microsystems, Wetzlar, Germany). An argon laser at 488 nm was used to detect fluorescence from the cytosolic calcium, and a 10× air objective was used to visualize the samples. For each sample, a movie was taken at a single focal plane for 2 hours at a rate of 1 frame / 15 seconds. Image analysis and data processing were performed in Matlab (The MathWorks, Inc., Natick, MA, USA).

### Quantifying single-cell Ca^2+^ oscillations

Each cell’s fluorescent intensity *I*(*t*) was obtained by averaging the pixel color values of 25 pixels around each cell’s center of mass as the cell fluoresced in the neighborhood of the edge; the distance of the center of mass from the wound edge was measured. The response curve *R*(*t*) was obtained by normalizing *I*(*t*) with a baseline intensity *I_r_* that was obtained by averaging each cell’s raw intensity over the last 5 minutes of the recording, which was used as a reference intensity due to the low value of the Ca^2+^ dye intensity that was caused by photobleaching. The response curve *R*(*t*) was then constructed by calculating *R*(*t*) = (*I*(*t*) − *I_r_*)/*I_r_*. The oscillations were next detected in Matlab using the function peakfinder.m (available through Matlab File Exchange), which identified local peaks in *R*(*t*). The peaks were defined as local maxima with values greater than zero and above the surrounding data by 0.5. If a cell’s response in time had two or more peaks, we referred to the cell as an oscillating cell (49).

### Traction force microscopy

Measurements of the traction exerted on the elastic substrates were done following established protocols (53, 54). Details are described in the supplementary material. Briefly, we recorded images of the cells and fluorescent beads following creation of the monolayer edge. After imaging, cells were removed by addition of trypsin. Substrate deformations below the cells were extracted from the image pairs with and without cells by particle image velocimetry. The deformation data was subsequently used to calculate the planar traction on the gel surface. This calculation was done in Fourier space, and consisted of inverting a Green’s function relating traction to displacement on a gel with a finite thickness (20, 55). The effect of measurement noise was suppressed by regularization of the traction magnitude (56).

## RESULTS

### Fibroblast cells exhibit calcium oscillations in the neighborhood of a wound edge

We image calcium dynamics in fibroblast monolayers in a region indicated by the red rectangle in figure 1(a). Monolayers have an areal density of >500 cells/mm^2^. Imaging is done before and after the removal of the PDMS strip. Figure 2 shows a typical example of Ca^2+^ dynamics in one cell. Prior to creation of the wound edge, most cells exhibit a flat Ca^2+^ temporal profile as shown in figure 2(a). After creation of an edge, most cells exhibit Ca^2+^ oscillations (figure 2(b)). Note that we are not able to monitor the cytosolic Ca^2+^ intensity for more than 2 hours due to the decaying intensity of the Fluo-4 Ca^2+^ dye caused by photobleaching; however, 2 hours provide sufficient time to identify a characteristic Ca^2+^ behavior in the wounded fibroblast monolayers. Culturing cells on substrates with different rigidity affects the calcium response. Cell monolayers on more rigid substrates show a higher number of oscillating cells following the creation of the wound (figure 2(c)). For a Young’s modulus *E* ~ 25 kPa, wounding causes a ~70% increase in the fraction of cells that exhibit Ca^2+^ oscillations.

**Figure 2.**
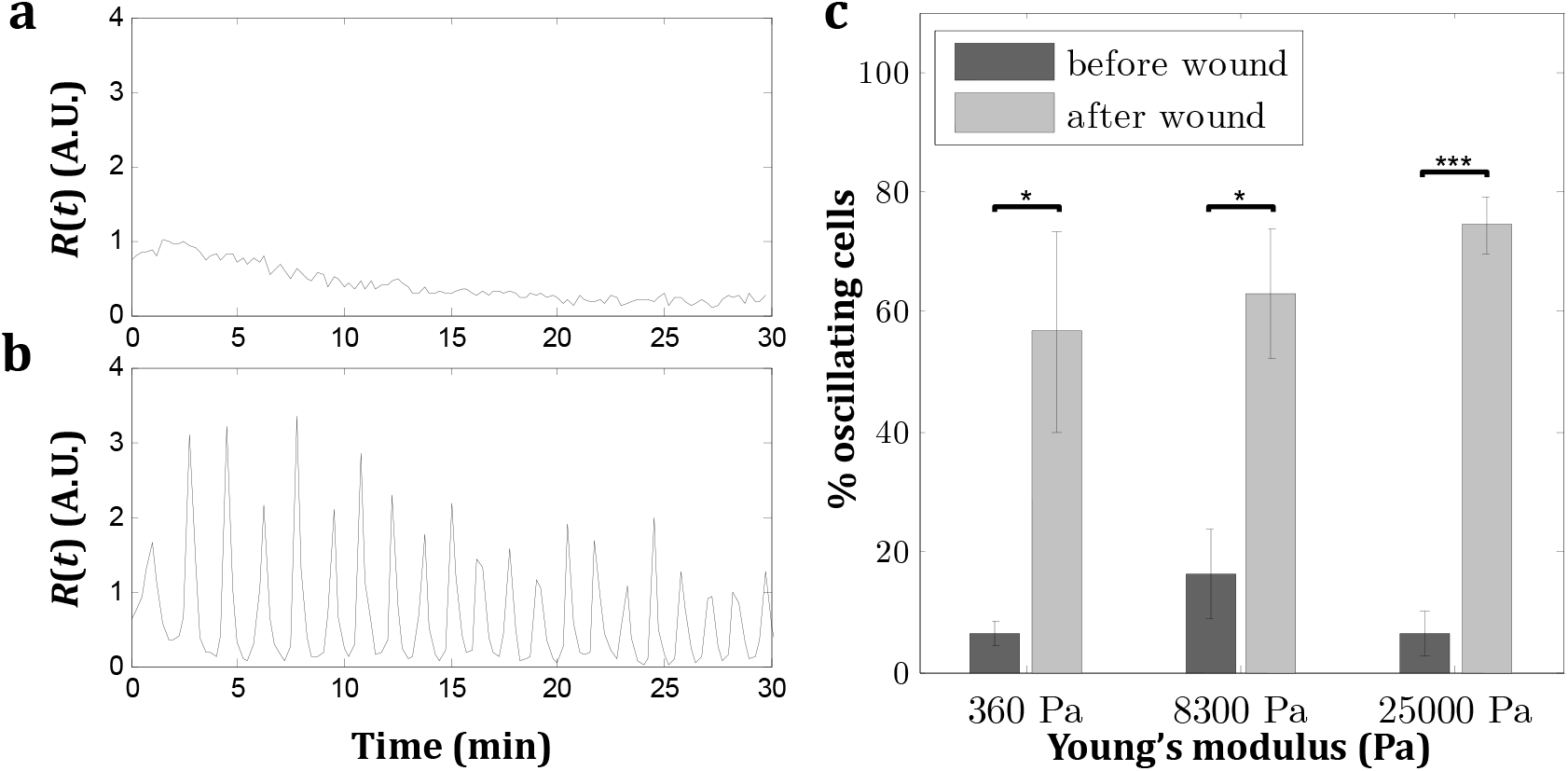
Calcium dynamics in fibroblast monolayers with areal density of >500 cells/mm^2^. (a) Typical Ca^2+^ profile in fibroblast monolayer on PA gel, *E* = 8300 Pa, before exposure to wound edge. (b) Typical Ca^2+^ profile in fibroblast monolayer on PA gel, *E* = 8300 Pa, after exposure to wound edge. (c) Significantly more cells show Ca^2+^ oscillations upon exposure to the wound edge following removal of the PDMS strip. Values are the average of three experiments and error bars are standard errors of the mean (\**p* ≤ 0.05, *\*\*\*p* ≤ 0.001).

### Wound-induced calcium oscillation period varies with the cell density

Previously, we had observed that calcium oscillations in fibroblast monolayers without a free edge can be stimulated by ATP in a cell-density dependent manner (49). To test if wound-induced oscillations are also density-dependent, we create cell monolayers of varying density (400-1200 cells/mm^2^) on PA gel, *E* = 8300 Pa (figure 3(a)), and quantified the Ca^2+^ oscillations within ~1 mm distance from the edge. The calcium oscillation period is defined as the time between successive maxima of the fluorescence intensity. We observe that the Ca^2+^ oscillation period varies with cell density, as shown in figure 3(b). For each sample, we observe a distribution of oscillation periods (figure 3(c)). By taking the mode of these distributions as the dominant oscillation period for a monolayer, we observe that the oscillation period for wounded fibroblast monolayers decreases with cell density (figure 3(d)). Thus, wound-induced calcium oscillations have a collective nature, as also confirmed by the increasing number of oscillating cells with increasing cell density for various substrates (figure S3 in the supplementary material). This collective calcium response is consistent with our previous finding where cells that are located closer to each other in a monolayer exhibit higher fraction of Ca^2+^-oscillating cells when excited by ATP (49).

**Figure 3.**
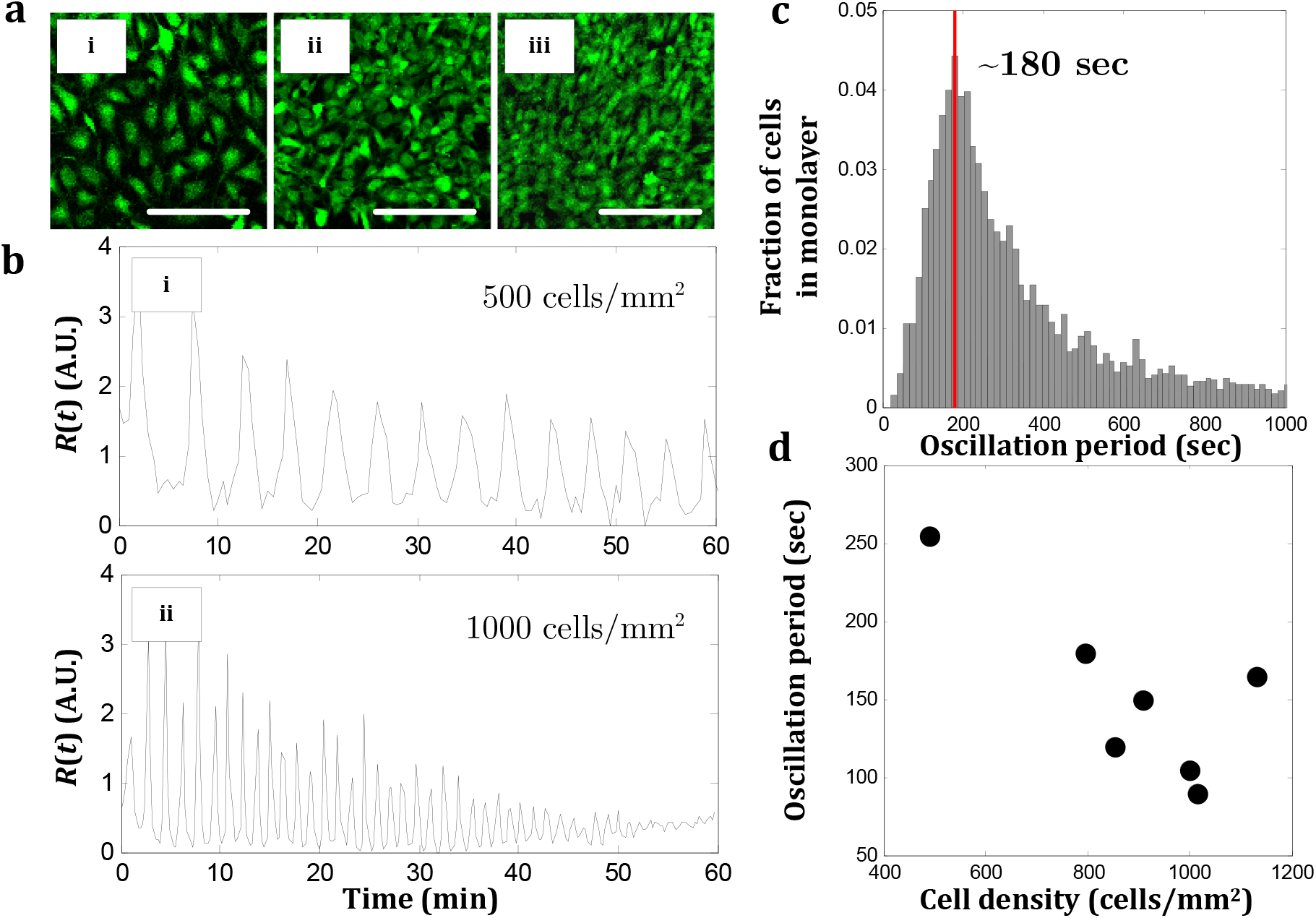
Effect of cell density on the period of wound-induced calcium oscillations. (a) Fibroblast cell monolayers on PA gels of various areal densities: (i) 500 cells/mm^2^, (ii) 800 cells/mm^2^, and (iii) 1000 cells/mm^2^. Scale bar: 200 μm. (b) Ca^2+^ oscillations in fibroblast cells on PA gel, *E* = 8300 Pa, following exposure to a free edge for monolayer density of: (i) 500 cells/mm^2^, and (ii) 1000 cells/mm^2^. (c) Distribution of Ca^2+^ oscillation periods in an intermediate density monolayer (800 cells/mm^2^) on PA gel, *E* = 8300 Pa. (d) Dependence of Ca^2+^ oscillation period on cell density, *E* = 8300 Pa.

The time intervals between successive calcium peaks vary considerably, which may be attributed to variations on the level of individual cells. However, the scatter of oscillation periods can also be caused by variability among different cells. Considering the data in figure 3(c), one may be tempted to describe calcium spiking of individual cells as a stochastic process where the time intervals between spikes follow a Poisson distribution. To test how cell-cell variations cause spread of the oscillation period, we follow Skupin et al. (57) and Thurley et al. (58) and calculate the average oscillation period together with the variance of the oscillation period for individual cells (figure S4(a-b) in the supplementary material). As shown earlier for other types of cells (58), we find a linear correlation between the average oscillation period and the standard deviation of the oscillation periods for fibroblasts. Thus, variability of oscillations on the single cell level is proportional to the period of oscillations.

This result is consistent with the interpretation that large oscillation periods allows many stochastic perturbations during each time interval between spikes. Note however that the linear correlation between standard deviation and average of the period has a finite offset and a slope that is smaller than unity (figure S4(b)). The linear correlation implies that oscillation periods on a single cell level are clearly not governed by a one-parameter Poisson process, which is also evident from observing the regularity of the oscillation signal in figure 3(b).

The cell-cell variability of oscillations can be gauged by observing the spread of average oscillation periods (figure S4(b) and S5 in the supplementary material). We observe average periods ranging between <100 seconds up to several 100 seconds. Thus, cell-cell variability is a major contributor to the uncertainty of the oscillation periods.

Note that the period of wound-induced Ca^2+^ oscillations can be significantly longer (average of ~250 sec for low cell density monolayers) than those of ATP-induced Ca^2+^ oscillations in an identical system (~100 sec) (49). When cultured on various substrates (*E* = 360 Pa − 10 GPa), the wound-induced Ca^2+^ oscillation period shows no general trend of dependence on the substrate Young’s modulus (figure S6 in the supplementary material). The absence of a systematic dependence of oscillation period on substrate rigidity is consistent with results from our previous studies where ATP-induced Ca^2+^ oscillations were investigated in a similar fibroblast monolayer system (49).

### Inhibition of myosin-II activity reduces wound-induced calcium oscillations

The creation of an edge in a cell monolayer is a strong mechanical perturbation. We hypothesized that the calcium response following this external perturbation also depends on the internal tensional state of the cells in the monolayer. To test this hypothesis, we treat high density cell monolayers (~1000 cells/mm^2^) on a PA gel, *E* = 25000 Pa with 50 μM blebbistatin (Sigma-Aldrich) for 30 minutes. Blebbistatin is a myosin II inhibitor that reduces cytoskeletal force generation, prevents the formation of circumferential actin bundles, and disrupts the coherence of the actin network in fibroblasts (59). Reduction of contractility by blebbistatin affects a range of mechanobiological systems that allow sensing of the extracellular environment, such as cadherin-based cell-cell adhesions and integrin-based adhesions to the extracellular substrate. In our experiments, we find that treatment with blebbistatin results in a decrease in the number of cells that exhibit Ca^2+^ oscillatory behavior (figure S7(a) in the supplementary material), which is consistent with our previously reported observations in a similar cell culture system stimulated by ATP (49). Thus, our results support the notion that myosin-based contractility influences intracellular Ca^2+^ dynamics. The interplay between the changes in cell-cell adhesion and cell-substrate adhesion due to blebbistatin treatment likely causes a slight increase in oscillation period (figure S7(b)).

### Localization of wound-induced calcium oscillations depends on substrate elasticity

To assess if the calcium response to wound creation is confined to the proximity of the wound, we quantify the occurrence of intracellular Ca^2+^ oscillations as a function of the distance from the wound edge. On rigid substrates such as glass, we observe that cells that show the most Ca^2+^ oscillation peaks are located away from the wound edge, as illustrated by the number of colored cells at ~700 μm away from the wound edge (white rectangle, figure 4(a)). Next, we calculate the sum of the number of Ca^2+^ oscillation at any given distance from the wound edge and ask how the oscillations are spatially distributed in the monolayers. We find that the spatial distribution of Ca^2+^ oscillations is a function of the substrate elasticity. For cells on soft substrates (PA gel, *E* = 360 Pa), most Ca^2+^ oscillations occur at ~0.05 mm distance from the wound edge (figure 4(b)), while on stiff substrates such as glass, most oscillations occur at up to ~0.85 mm away from the wound edge (figure 4(c)). By comparing the average of the modes of the Ca^2+^ oscillation distributions on various substrates, we show that decreasing substrate rigidity generally leads to closer localization of calcium activity to the wound (figure 4(d)). This average distance of the maximum Ca^2+^ activity from the wound edge for the various substrates ranges from ~10 to 60 times the cell diameter. It is also interesting to note that while it is known that substrate stiffness changes the cell spreading area for individual cells (53), we observe no significant difference in spreading area of our wounded 3T3 monolayers (figure S8 in the supplementary material), therefore the area of cell-cell contacts is also likely very similar on different substrates.

**Figure 4.**
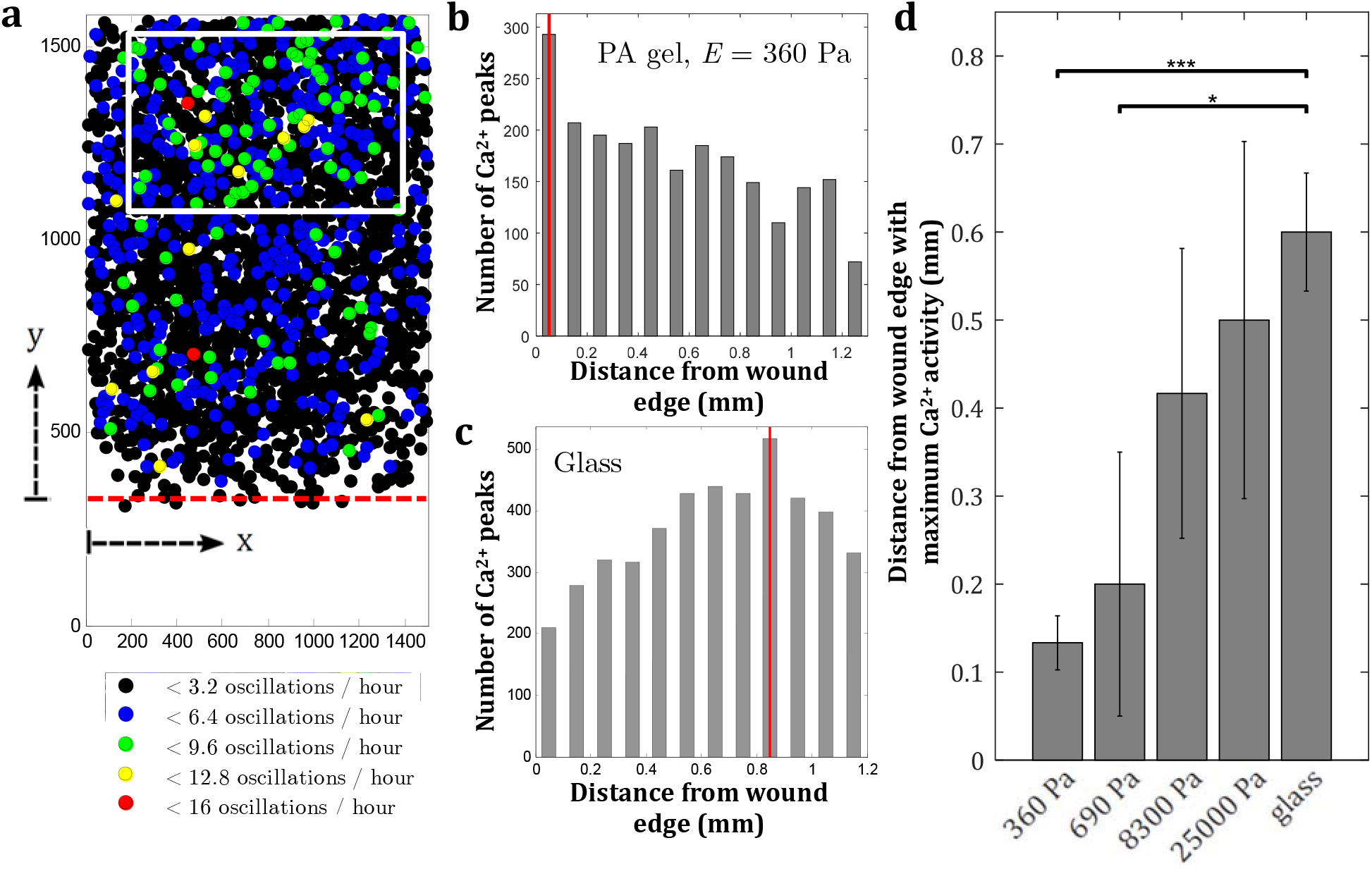
Spatial behavior of calcium oscillations in a wounded monolayer depends on substrate elasticity. (a) Spatial distribution of the cells exhibiting Ca^2+^ oscillations on glass. The *y*-position corresponds to the distance from the wound edge. (b–c) Examples of the distribution of the total number of Ca^2+^ peaks as a function of distance from the edge for cells cultured on PA gel, *E* = 360 Pa (b) and on glass (c). Maximum Ca^2+^ activity occurs around 0.05 mm from the edge for soft PA gel (b) and 0.85 mm from the edge for glass (c). (d) For all substrates, *E* = 360 Pa − *O*(10 GPa), the average distance of maximum Ca^2+^ activity from the wound edge increases with substrate stiffness, up to ~60 cell diameters. Values are the averages of six experiments and error bars are standard errors of the mean (\**p* ≤ 0.05, *p*\*\*\*≤0.001).

Measurement of a higher number of calcium peaks can result from more oscillating cells, from cells oscillating more persistently, or from higher oscillation frequency. Defining oscillating cells as those having at least two calcium spikes, we plot the distance from the wound edge with the maximum number of oscillating cells for the various substrates (figure S9 in the supplementary material). The average distance from the wound edge with most oscillating cells increases with substrate stiffness. Although the trend seems weak, this finding follows the same trend as figure 4(d), where in softer substrates the highest number of Ca^2+^ oscillations are seen closer to the edge. For stiff substrates (*E* = 25 kPa and glass), the distance with most oscillating cells is roughly the same as the distance with most Ca^2+^ oscillations, around 0.5-0.6 mm (figure 4(d), figure S9). However, for the softer substrates, these distances are different from each other, i.e. the distance with most oscillating cells is greater than distance with most Ca^2+^ oscillations, suggesting there are cells on these softer substrates (*E =* 360 − 8300 Pa) closer to the edge that oscillate more persistently after wound creation, giving rise to high number of Ca^2+^ oscillations at a shorter distance from the wound edge.

To next clarify whether the calcium oscillation periods vary spatially, we also examine the dependence of the oscillation periods on the distance *y* from the wound edge. As shown in figure S5, the average distribution of oscillation periods does not depend in a systematic way on *y*. Therefore, the distribution of calcium oscillations in figure 4(d) does not result from oscillation period variations. To finally test if the regularity of oscillations possibly depends on the spatial distance from the edge, we consider the standard deviation divided by the average of the periods on the level of individual cells. As demonstrated in figure S4(c-e), no preference for regular oscillations is observed close to the edge.

### Spatial localization of cell-substrate traction correlates with calcium oscillation

Cells feel the extracellular substrate rigidity by exerting traction. Since the substrate rigidity affects localization of calcium oscillations, we concluded that the cell-substrate traction likely plays an important role. To measure the distribution of cell-substrate forces after creation of a wound edge, we employ a traction force microscopy method for cell sheets, as pioneered in Del Alamo et al. and Trepat et al. (20, 55). Traction forces are measured after the removal of the PDMS strip. On a soft PA gel with Young’s modulus *E* = 690 Pa, high traction magnitude of ~40 Pa is observed locally near the monolayer edge (figure 5(a,b)). To compare data from different experiments, we average the traction *t_xy_* over the *x*-coordinate, which is approximately parallel to the colony edge, as 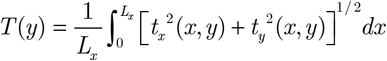. Here, *L_x_* is the width of the image. Figure 5(a-b) shows that the traction distributions measured in different samples on soft substrates are consistent with the notion that the most traction occurs close to the edge. At distances larger than ~200 μm away from the edge, the traction is much smaller than at the edge but does not completely vanish. The traction distribution that we observe for fibroblasts on soft substrates is in qualitative agreement with the traction distribution reported for migrating epithelial cell sheets (20).

**Figure 5.**
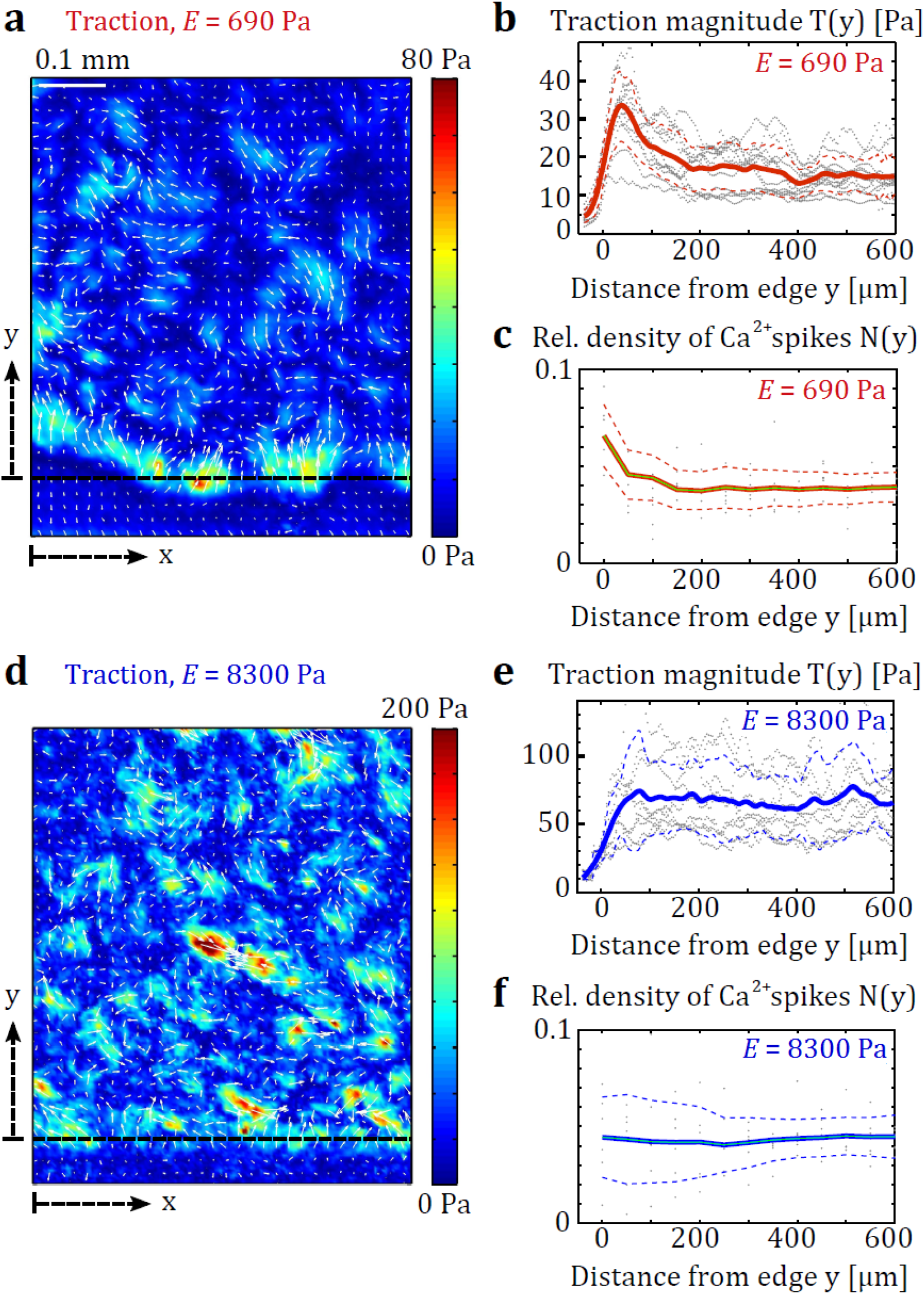
Spatial distribution of traction in wounded fibroblast monolayers varies with substrate elasticity. (a,d) Map of traction distribution in wounded fibroblast monolayers on PA gels. Quivers indicate direction of traction. (b,e) Traction magnitude profile as a function of distance from the wound edge. The origin of the *y*-coordinate is approximately located at the wound edge. Non-vanishing traction below *y* = 0 results from limited spatial traction resolution, noise, and the wavy shape of the edge. Gray points: original data, lines: average over different experiments, dashed lines: standard deviation of data. At least five experiments are repeated for each condition. (c,f) Relative density profile of Ca^2+^ oscillations. Note that the distributions of traction magnitude and calcium oscillations co-localize on both substrate rigidities.

On stiffer PA gels, however, the monolayers exert different traction fields. For a Young’s modulus *E* larger than about 2 kPa, traction occurs no longer preferentially at the edge (figure S2 in the supplementary material). On rigid substrates with *E* = 8300 Pa, small areas with high traction occur everywhere in the monolayer (figure 5(d)). Averaging the traction along the *x*-coordinate leads to a statistical distribution of traction that is almost constant in space, where magnitudes of ~50 Pa are measured up to ~600 μm away from the edge of the wounded monolayer (figure 5(e)). Also, we find that the traction magnitude is substantially larger on stiff substrates than it is on soft substrates (figure 5, figure S2 in the supplementary material).

Since the absolute number of calcium oscillations varies from experiment to experiment, comparison of the experimental conditions is facilitated by normalizing the number of locally occurring oscillations by the overall number of oscillations in one experiment. We denote by *n* = *n* (*x*, *y)* the overall number of calcium oscillations occurring during the imaging at the point (*x*, *y*). Then, the relative density of oscillations occurring in a monolayer with size *L_x_* × *L_y_* is defined as:

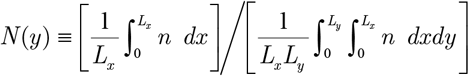

As seen in figures 5(b-c,e-f), the relative density of calcium oscillations co-localizes with the distribution of traction magnitude measured on different substrate rigidities. Note that this data does not mean that traction magnitude and number of calcium oscillations co-localize precisely on the *x*-*y* plane. Rather, both quantities depend on average similarly on the distance from the edge; i.e. high traction and high relative oscillation density are observed locally near the monolayer edge for *E* = 690 Pa, and spatially distributed traction and spatially distributed relative oscillation density are observed for cells on stiff PA gel, *E* = 8300 Pa.

The local imbalance of substrate traction can be used to estimate the intercellular stress between cells in the monolayer. The intercellular stress σ_*yy*_ in the *y*-direction is proportional to the integral of the traction component *t_γ_* as 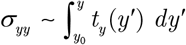. In contrast to traction magnitude, the stress σ_*yy*_ qualitatively increases with distance from the edge on soft substrates (figure S2(d) in the supplementary material). This trend of increasing stress does not match the distribution of calcium oscillations. Therefore, localization of wound-induced calcium oscillations in fibroblasts is primarily linked to cell-substrate traction, while intercellular stress likely plays only a secondary role.

## DISCUSSION

Calcium is a paradigmatic regulator of cellular activity. For fibroblasts, it has been observed that the cytosolic level of calcium changes after chemical or mechanical stimulation (49, 60–65). Here, we ask if fibroblasts can detect and localize tissue wounds through calcium signaling. We find that creation of an edge in a fibroblast monolayer can indeed trigger oscillations of the cytosolic calcium in the majority of the monolayer cells. Wound-induced oscillations are much more persistent than the previously reported oscillations following chemical stimulation by ATP (49). Measurable oscillations can last for > 2 hours after wounding, which is comparable to the time it takes to start cellular reorientation and migration. We conclude that similar timescales of calcium excitation and cell organization could in principle allow a causal connection between calcium dynamics and mechanical reorganization following wound creation.

We next vary the density of cells in the monolayer to see if calcium oscillations are affected by cellular crosstalk. Increase of cell density leads to higher oscillation frequency and a higher number of oscillating cells. This finding suggests that wound-induced calcium oscillations, similar to agonist-induced oscillations, are a collective effect where the calcium signal is propagated via gap junctions from cell to cell (61, 66). Gap junctions in 3T3 cells were identified as crucial in preserving the spatial pattern of Ca^2+^ communication during the initial stages of collective chemosensing (59), and its inhibition by palmitoleic acid changes the dynamics of the Ca^2+^ oscillations (49). The importance of short-range gap junction communication in regulating collective Ca^2+^ dynamics is also supported by nearest neighbor cross-correlation analysis of the Ca^2+^ responses in our wounded 3T3 systems, where synchronization events are observed as indicated by the positive correlation peak at zero time delay (figure S10 in the supplementary material). At regions of 3T3 cell-cell contacts, cadherins are present and are colocalized with connexin43 (C×43 α1) gap junction protein, where knockdown of either protein results in reduced gap junctional communication and inhibition of cell motility (67). The formation of gap junction, adherens junction, and the association between the two depend on the intracellular coassembly of connexin and cadherin, thus confirming the relevance of investigating Ca^2+^ communication through gap junctions and force transmission through cell-cell junctions in our system of wounded fibroblast monolayer.

Our current and previous studies suggest that dynamic coupling of cells through gap junction is the dominant mode of Ca^2+^ communication following wounding and ATP excitation in 3T3 cell monolayers (49, 59). It is also known that gap junctions can close upon elevated intracellular Ca^2+^ level (68). Despite this evidence for the role of gap junctions, the observed oscillations can also occur without this short-ranged type of interactions, as seen in 3T3 cells embedded in gels (49), suggesting the presence of diffusive paracrine signaling in this system. The coupling between gap junction dynamics, paracrine signaling, wound-induced Ca^2+^ release, and Ca^2+^-induced Ca^2+^ release can allow the oscillations to synchronize, therefore also contributing to the large variability in the oscillation period.

Regarding Ca^2+^ oscillations in 3T3 fibroblast cells, involvements of Ca^2+^ stores and second messenger IP3 has previously been investigated quite extensively. In single cells, inhibition of sarcoplasmic-endoplasmic Ca^2+^-ATPase (SERCA) using thapsigargin results in the depletion of Ca^2+^ from Ca^2+^ storage, triggering Ca^2+^ influx (69–73). Mechanically, an intact cytoskeleton is required for agonist-induced Ca^2+^ signaling, not through lower capacitative calcium entry, impaired IP3 receptor function, or diminished phospholipase C (PLC) activity, but through alteration of spatial relationship between PLC and IP3 receptors (74). The severe rearrangement of endoplasmic reticulum (ER) membranes, the site for PLC generation of IP3, also confirmed that such spatial change could impair PLC-dependent calcium signaling.

Earlier studies demonstrate that the tensile state of intracellular actin controls Ca^2+^ activities in various cell types (75–77). Moreover, opening of mechanosensitive Ca^2+^ channels, which allows entry of extracellular Ca^2+^, are regulated by stress fiber formation and contraction (18, 78). To test if our wound-induced calcium oscillations also depend on cytoskeletal tension, we partially inhibit cell contractility using blebbistatin. After disruption of contractility, fewer cells show calcium oscillations. This finding directly confirms the coupling between actomyosin activity and calcium response in wounded fibroblast monolayers.

Cytoskeletal tension of adherent cells is directly transmitted to the extracellular substrate via integrin-based adhesions. Since tension affects calcium oscillations, we examine how the extracellular matrix rigidity affects oscillations. To this end, cell monolayers are grown on substrates with different elastic modulus. We find that an increased rigidity of the substrate leads to an increase of the number of oscillating cells. However, the oscillation frequency varies non-monotonically with substrate rigidity. This observation suggests that mechanonsensitivity of fibroblast calcium oscillations is more systematically encoded in excitability than in the oscillation frequency. Consequently, we next focus on the spatial distribution of the number of calcium oscillations to determine if cells react locally to the creation of a wound edge. We find that generally, cells throughout the observed region in the monolayer may show calcium oscillations after wound edge creation. However, with decreasing substrate rigidity, the mode of the spatial oscillation distribution moves closer to the monolayer edge. For soft substrates with *E* < ~2 kPa, we observe a pronounced localization of the number of Ca^2+^ oscillations within a region on the order of 100 μm around the edge. Thus, calcium oscillations co-localize with the wound edge on soft substrates.

To establish a possible link between mechanosensitivity of calcium oscillations and edge-localization on soft substrates, we next investigate the cell-substrate traction of our fibroblast monolayers. For epithelial cell monolayers on a soft substrate (*E* = 1.3 kPa), it has been reported that the traction normal to the edge is highest near the edge and decays to a non-zero average value away from the edge (20, 79). Although epithelial cell sheets undergo efficient durotaxis (80), their collective response to substrate stiffness has been reported to be less pronounced than for individual cells (20). In contrast, endothelial cell sheets (HUVEC) respond to a higher substrate rigidity by producing higher traction (81). Likewise, we find for fibroblast monolayers an increased traction magnitude with higher substrate rigidity. However, we find that substrate rigidity not only affects traction magnitude, but also traction distribution for our system. For substrates with *E* < ~2 kPa, we observe a strong localization of absolute traction magnitude to the newly created edge of the cell monolayer. On more rigid substrates, traction magnitude is statistically uniform in the monolayer.

The rigidity-dependent edge localization of traction could be plausibly explained by a change of the force balance among cadherin-based intercellular connection and integrin-based cell-matrix connections. On soft substrates, we assume that the forces are preferentially transmitted via cell-cell adhesions, which leads to a traction build-up at the monolayer edge where no neighboring cells are present. On stiff substrates, the fraction of force transmitted by cells directly to the substrate is larger, leading to less force at the monolayer edge. This interpretation of our data is consistent with the observation that individual fibroblasts require a substrate rigidity above around *E* ~ 8-9 kPa to form pronounced adhesion sites and actin stress fibers, while cells in dense collections can maintain the same morphology on soft and stiff substrates (53).

In this study on calcium dynamics, we find that upon the creation of a wound, cell-substrate traction and calcium oscillations both localize close to the wound if the substrate is soft, *E* < 2 kPa. Our results suggest that the higher number of calcium spikes at the edge results from a local buildup of traction which in turn is determined by the balance of cell-substrate and cell-cell forces in the monolayer. It has previously been shown that spontaneous calcium oscillations are mainly transmitted via gap junctions among fibroblasts, while inhibition of OB-cadherins and N-cadherins does not affect oscillation periods significantly (78). Based on our own results and these earlier studies, we believe that intercellular mechanical stress does not affect calcium oscillations in a direct way, but likely affects the overall excitability of the cell through an interplay with substrate forces. Gap-junctional connections, in contrast, likely directly excite oscillations (59).

Further studies are needed to separate the effects of cell-substrate forces, intercellular forces, and intercellular signaling, which are all necessary for Ca^2+^ oscillations in fibroblasts. In future research, careful titration of pharmacological inhibitors will hopefully produce a more detailed picture of the mechanisms underlying calcium oscillations after wounding that was reported here. The observed spatial correlation between calcium oscillations and traction in our study suggests a causal link between both, leading us to speculate that the described mechanism of collective edge localization by calcium oscillations may play an unknown physiological role.

## CONCLUSIONS

We have demonstrated that calcium oscillations are modulated by forces associated with wound creation in fibroblast cell monolayers. Our observations lead us to conclude that collective calcium oscillatory behavior can allow localization of wound edges in a substrate-dependent fashion. The study highlights a potential significance of wound-induced calcium dynamics in diverse microenvironments.

## ACKNOWLEDGEMENTS

We thank members of the Stone laboratory for discussions and advice. We particularly acknowledge Dr. Antonio Perazzo for the help with rheometry. B.S. was supported by a postdoctoral fellowship from the DAAD.

## REFERENCES

1. Martin P. Wound healing––aiming for perfect skin regeneration. Science. 1997 Apr 4;276(5309):75–81.

2. Fenteany G, Janmey PA, Stossel TP. Signaling pathways and cell mechanics involved in wound closure by epithelial cell sheets. Curr Biol. 2000 Jul 13;10(14):831–8.

3. Friedl P, Gilmour D. Collective cell migration in morphogenesis, regeneration and cancer. Nat Rev Mol Cell Biol. 2009 Jul;10(7):445–57.

4. Ladoux B, Anon E, Lambert M, Rabodzey A, Hersen P, Buguin A, et al. Strength Dependence of Cadherin-Mediated Adhesions. Biophys J. 2010 Feb 17;98(4):534–42.

5. Huttenlocher A, Horwitz AR. Integrins in Cell Migration. Csh Perspect Biol. 2011 Sep;3(9).

6. Schiller HB, Fassler R. Mechanosensitivity and compositional dynamics of cell-matrix adhesions. Embo Rep. 2013 Jun;14(6):509–19.

7. Geiger B, Bershadsky A, Pankov R, Yamada KM. Transmembrane crosstalk between the extracellular matrix––cytoskeleton crosstalk. Nat Rev Mol Cell Bio. 2001 Nov;2(11):793–805.

8. Borghi N, Lowndes M, Maruthamuthu V, Gardel ML, Nelson WJ. Regulation of cell motile behavior by crosstalk between cadherin and integrin-mediated adhesions. P Natl Acad Sci USA. 2010 Jul 27;107(30):13324–9.

9. Tsai J, Kam L. Rigidity-Dependent Cross Talk between Integrin and Cadherin Signaling. Biophys J. 2009 Mar 18;96(6):L39–L41.

10. McCain ML, Lee H, Aratyn-Schaus Y, Kleber AG, Parker KK. Cooperative coupling of cell-matrix and cell-cell adhesions in cardiac muscle. P Natl Acad Sci USA. 2012 Jun 19;109(25):9881–6.

11. Martinez-Rico C, Pincet F, Thiery JP, Dufour S. Integrins stimulate E-cadherin-mediated intercellular adhesion by regulating Src-kinase activation and actomyosin contractility. Journal of Cell Science. 2010 Mar 1;123(5):712–22.

12. Weber GF, Bjerke MA, DeSimone DW. Integrins and cadherins join forces to form adhesive networks. Journal of Cell Science. 2011 Apr 15;124(8):1183–93.

13. Rottner K, Hall A, Small JV. Interplay between Rac and Rho in the control of substrate contact dynamics. Curr Biol. 1999 Jun 17;9(12):640–8.

14. Burridge K, Wennerberg K. Rho and Rac take center stage. Cell. 2004 Jan 23;116(2):167–79.

15. Machacek M, Hodgson L, Welch C, Elliott H, Pertz O, Nalbant P, et al. Coordination of Rho GTPase activities during cell protrusion. Nature. 2009 Sep 3;461(7260):99–103.

16. Omelchenko T, Vasiliev JM, Gelfand IM, Feder HH, Bonder EM. Rho-dependent formation of epithelial "leader" cells during wound healing. Proc Natl Acad Sci U S A. 2003 Sep 16;100(19):10788–93.

17. Howe AK. Cross-talk between calcium and protein kinase A in the regulation of cell migration. Curr Opin Cell Biol. 2011 Oct;23(5):554–61.

18. Hayakawa K, Tatsumi H, Sokabe M. Actin stress fibers transmit and focus force to activate mechanosensitive channels. J Cell Sci. 2008 Feb 15;121(Pt 4):496–503.

19. Poujade M, Grasland-Mongrain E, Hertzog A, Jouanneau J, Chavrier P, Ladoux B, et al. Collective migration of an epithelial monolayer in response to a model wound. Proc Natl Acad Sci U S A. 2007 Oct 9;104(41):15988–93.

20. Trepat X, Wasserman MR, Angelini TE, Millet E, Weitz DA, Butler JP, et al. Physical forces during collective cell migration. Nat Phys. 2009 Jun;5(6):426–30.

21. Tambe DT, Hardin CC, Angelini TE, Rajendran K, Park CY, Serra-Picamal X, et al. Collective cell guidance by cooperative intercellular forces. Nature Materials. 2011 Jun;10(6):469–75.

22. Maruthamuthu V, Sabass B, Schwarz US, Gardel ML. Cell-ECM traction force modulates endogenous tension at cell-cell contacts. P Natl Acad Sci USA. 2011 Mar 22;108(12):4708–13.

23. Mertz AF, Che YL, Banerjee S, Goldstein JM, Rosowski KA, Revilla SF, et al. Cadherin-based intercellular adhesions organize epithelial cell-matrix traction forces. P Natl Acad Sci USA. 2013 Jan 15;110(3):842–7.

24. Kalluri R, Weinberg RA. The basics of epithelial-mesenchymal transition. J Clin Invest. 2009 Jun;119(6):1420–8.

25. Evans JH, Falke JJ. Ca^2+^ influx is an essential component of the positive-feed back loop that maintains leading-edge structure and activity in macrophages. Proceedings of the National Academy of Sciences of the United States of America. 2007 Oct 9;104(41):16176–81.

26. Wei CL, Wang XH, Chen M, Ouyang K, Song LS, Cheng HP. Calcium flickers steer cell migration. Nature. 2009 Feb 12;457(7231):901–5.

27. Gough AH, Taylor DL. Fluorescence Anisotropy Imaging Microscopy Maps Calmodulin-Binding during Cellular Contraction and Locomotion. J Cell Biol. 1993 Jun;121(5):1095–107.

28. Harris AS, Morrow JS. Calmodulin and Calcium-Dependent Protease-I Coordinately Regulate the Interaction of Fodrin with Actin. P Natl Acad Sci USA. 1990 Apr;87(8):3009–13.

29. Hartwig JH, Thelen M, Rosen A, Janmey PA, Nairn AC, Aderem A. Marcks Is an Actin Filament Cross-Linking Protein Regulated by Protein-Kinase-C and Calcium Calmodulin. Nature. 1992 Apr 16;356(6370):618–22.

30. Cunningham CC, Stossel TP, Kwiatkowski DJ. Enhanced Motility in Nih-3t3 Fibroblasts That Overexpress Gelsolin. Science. 1991 Mar 8;251(4998):1233–6.

31. Hendey B, Klee CB, Maxfield FR. Inhibition of Neutrophil Chemokinesis on Vitronectin by Inhibitors of Calcineurin. Science. 1992 Oct 9;258(5080):296–9.

32. Zhang X, Chattopadhyay A, Ji QS, Owen JD, Ruest PJ, Carpenter G, et al. Focal adhesion kinase promotes phospholipase C-gamma l activity. P Natl Acad Sci USA. 1999 Aug 3;96(16):9021–6.

33. Renshaw MW, Ren XD, Schwartz MA. Growth factor activation of MAP kinase requires cell adhesion. Embo J. 1997 Sep 15;16(18):5592–9.

34. Renshaw MW, Price LS, Schwartz MA. Focal adhesion kinase mediates the integrin signaling requirement for growth factor activation of MAP kinase. J Cell Biol. 1999 Nov 1;147(3):611–8.

35. Giannone G, Dubin-Thaler BJ, Rossier O, Cai Y, Chaga O, Jiang G, et al. Lamellipodial actin mechanically links myosin activity with adhesion-site formation. Cell. 2007 Feb 9;128(3):561–75.

36. Tsai FC, Meyer T. Ca^2+^ Pulses Control Local Cycles of Lamellipodia Retraction and Adhesion along the Front of Migrating Cells. Curr Biol. 2012 May 8;22(9):837–42.

37. Franco SJ, Rodgers MA, Perrin BJ, Han JW, Bennin DA, Critchley DR, et al. Calpain-mediated proteolysis of talin regulates adhesion dynamics. Nat Cell Biol. 2004 Oct;6(10):977–+.

38. Gomez TM, Snow DM, Letourneau PC. Characterization of Spontaneous Calcium Transients in Nerve Growth Cones and Their Effect on Growth Cone Migration. Neuron. 1995 Jun;14(6):1233–46.

39. Robles E, Huttenlocher A, Gomez TM. Filopodial calcium transients regulate growth cone motility and guidance through local activation of calpain. Neuron. 2003 May 22;38(4):597–609.

40. Komuro H, Rakic P. Intracellular Ca^2+^ fluctuations modulate the rate of neuronal migration. Neuron. 1996 Aug;17(2):275–85.

41. Mandeville JTH, Ghosh RN, Maxfield FR. Intracellular Calcium Levels Correlate with Speed and Persistent Forward Motion in Migrating Neutrophils. Biophys J. 1995 Apr;68(4):1207–17.

42. Tsai FC, Seki A, Yang HW, Hayer A, Carrasco S, Malmersjo S, et al. A polarized Ca^2+^, diacylglycerol and STIM1 signalling system regulates directed cell migration. Nat Cell Biol. 2014 Feb;16(2):133–44.

43. Pelham RJ, Wang YL. Cell locomotion and focal adhesions are regulated by substrate flexibility. P Natl Acad Sci USA. 1997 Dec 9;94(25):13661–5.

44. Wells RG. The role of matrix stiffness in regulating cell behavior. Hepatology. 2008 Apr;47(4):1394–400.

45. Discher DE, Janmey P, Wang YL. Tissue cells feel and respond to the stiffness of their substrate. Science. 2005 Nov 18;310(5751):1139–43.

46. Chen CS, Mrksich M, Huang S, Whitesides GM, Ingber DE. Geometric control of cell life and death. Science. 1997 May 30;276(5317):1425–8.

47. Vogel V, Sheetz M. Local force and geometry sensing regulate cell functions. Nat Rev Mol Cell Bio. 2006 Apr;7(4):265–75.

48. Lo CM, Wang HB, Dembo M, Wang YL. Cell movement is guided by the rigidity of the substrate. Biophysical Journal. 2000 Jul;79(1):144–52.

49. Lembong J, Sabass B, Sun B, Rogers ME, Stone HA. Mechanics regulates ATP-stimulated collective calcium response in fibroblast cells. Journal of the Royal Society, Interface / the Royal Society. 2015 Jul 6;12(108):20150140.

50. Godbout C, Castella LF, Smith EA, Talele N, Chow ML, Garonna A, et al. The Mechanical Environment Modulates Intracellular Calcium Oscillation Activities of Myofibroblasts. PLoS ONE. 2013 May 14;8(5).

51. Nemir S, West JL. Synthetic Materials in the Study of Cell Response to Substrate Rigidity. Ann Biomed Eng. 2010 Jan;38(1):2–20.

52. Chen EJ, Novakofski J, Jenkins WK, OBrien WD. Young’s modulus measurements of soft tissues with application to elasticity imaging. Ieee T Ultrason Ferr. 1996 Jan;43(1):191–4.

53. Yeung T, Georges PC, Flanagan LA, Marg B, Ortiz M, Funaki M, et al. Effects of substrate stiffness on cell morphology, cytoskeletal structure, and adhesion. Cell Motil Cytoskeleton. 2005 Jan;60(1):24–34.

54. Plotnikov SV, Sabass B, Schwarz US, Waterman CM. High-resolution traction force microscopy. Methods Cell Biol. 2014;123:367–94.

55. del Alamo JC, Meili R, Alonso-Latorre B, Rodriguez-Rodriguez J, Aliseda A, Firtel RA, et al. Spatio-temporal analysis of eukaryotic cell motility by improved force cytometry. P Natl Acad Sci USA. 2007 Aug 14;104(33):13343–8.

56. Sabass B, Gardel ML, Waterman CM, Schwarz US. High resolution traction force microscopy based on experimental and computational advances. Biophys J. 2008 Jan 1;94(1):207–20.

57. Skupin A, Kettenmann H, Winkler U, Wartenberg M, Sauer H, Tovey SC, et al. How does intracellular Ca^2+^ oscillate: By chance or by the clock? Biophys J. 2008 Mar 15;94(6):2404–11.

58. Thurley K, Tovey SC, Moenke G, Prince VL, Meena A, Thomas AP, et al. Reliable Encoding of Stimulus Intensities Within Random Sequences of Intracellular Ca^2+^ Spikes. Sci Signal. 2014 Jun 24;7(331).

59. Cai YF, Rossier O, Gauthier NC, Biais N, Fardin MA, Zhang X, et al. Cytoskeletal coherence requires myosin-IIA contractility. Journal of Cell Science. 2010 Feb 1;123(3):413–23.

60. Gonzalez FA, Rozengurt E, Heppel LA. Extracellular ATP induces the release of calcium from intracellular stores without the activation of protein kinase C in Swiss 3T6 mouse fibroblasts. Proc Natl Acad Sci U S A. 1989 Jun;86(12):4530–4.

61. Sun B, Lembong J, Normand V, Rogers M, Stone HA. Spatial-temporal dynamics of collective chemosensing. Proc Natl Acad Sci U S A. 2012 May 15;109(20):7753–8.

62. Pinheiro AR, Paramos-de-Carvalho D, Certal M, Costa MA, Costa C, Magalhaes-Cardoso MT, et al. Histamine induces ATP release from human subcutaneous fibroblasts, via pannexin-1 hemichannels, leading to Ca^2+^ mobilization and cell proliferation. J Biol Chem. 2013 Sep 20;288(38):27571–83.

63. Arora PD, Bibby KJ, McCulloch CA. Slow oscillations of free intracellular calcium ion concentration in human fibroblasts responding to mechanical stretch. J Cell Physiol. 1994 Nov;161(2):187–200.

64. Munevar S, Wang YL, Dembo M. Regulation of mechanical interactions between fibroblasts and the substratum by stretch-activated Ca^2+^ entry. Journal of Cell Science. 2004 Jan 1;117(Pt 1):85–92.

65. Furuya K, Furuya S, Yamagishi S. Intracellular calcium responses and shape conversions induced by endothelin in cultured subepithelial fibroblasts of rat duodenal villi. Pflugers Arch. 1994 Sep;428(2):97–104.

66. Sanderson MJ, Charles AC, Boitano S, Dirksen ER. Mechanisms and function of intercellular calcium signaling. Mol Cell Endocrinol. 1994 Jan;98(2):173–87.

67. Wei CJ, Francis R, Xu X, Lo CW. Connexin43 associated with an N-cadherin-containing multiprotein complex is required for gap junction formation in NIH3T3 cells. J Biol Chem. 2005 May 20;280(20):19925–36.

68. Alberts B, Johnson A, Lewis J, Morgan D, Raff M, Roberts K, et al. Molecular biology of the cell. 6 ed. New York: Garland Science; 2014.

69. Putney JW, Jr. A model for receptor-regulated calcium entry. Cell Calcium. 1986 Feb;7(1):1–12.

70. Polverino AJ, Hughes BP, Barritt GJ. Inhibition of Ca^2+^ inflow causes an abrupt cessation of growth-factor-induced repetitive free Ca^2+^ transients in single NIH-3T3 cells. Biochem J. 1991 Sep 15;278(Pt 3):849–55.

71. Ribeiro CM, Putney JW. Differential effects of protein kinase C activation on calcium storage and capacitative calcium entry in NIH 3T3 cells. J Biol Chem. 1996 Aug 30;271(35):21522–8.

72. Razani-Boroujerdi S, Partridge LD, Sopori ML. Intracellular calcium signaling induced by thapsigargin in excitable and inexcitable cells. Cell Calcium. 1994 Dec;16(6):467–74.

73. Miyakawa T, Kojima M, Ui M. Differential routes of Ca^2+^ influx in Swiss 3T3 fibroblasts in response to receptor stimulation. Biochem J. 1998 Jan 01;329(Pt 1):107–14.

74. Ribeiro CM, Reece J, Putney JW, Jr. Role of the cytoskeleton in calcium signaling in NIH 3T3 cells. An intact cytoskeleton is required for agonist-induced [Ca^2+^]i signaling, but not for capacitative calcium entry. J Biol Chem. 1997 Oct 17;272(42):26555–61.

75. Castella LF, Buscemi L, Godbout C, Meister JJ, Hinz B. A new lock-step mechanism of matrix remodelling based on subcellular contractile events. Journal of Cell Science. 2010 May 15;123(Pt 10):1751–60.

76. Dandekar SN, Park JS, Peng GE, Onuffer JJ, Lim WA, Weiner OD. Actin dynamics rapidly reset chemoattractant receptor sensitivity following adaptation in neutrophils. Philos T R Soc B. 2013 Nov 5;368(1629).

77. Holle AW, Engler AJ. More than a feeling: discovering, understanding, and influencing mechanosensing pathways. Curr Opin Biotech. 2011 Oct;22(5):648–54.

78. Follonier L, Schaub S, Meister JJ, Hinz B. Myofibroblast communication is controlled by intercellular mechanical coupling. J Cell Sci. 2008 Oct 15;121(20):3305–16.

79. Angelini TE, Hannezo E, Trepat X, Fredberg JJ, Weitz DA. Cell Migration Driven by Cooperative Substrate Deformation Patterns. Physical Review Letters. 2010 Apr 23;104(16).

80. Sunyer R, Conte V, Escribano J, Elosegui-Artola A, Labernadie A, Valon L, et al. Collective cell durotaxis emerges from long-range intercellular force transmission. Science. 2016 Sep 9;353(6304):1157–61.

81. Krishnan R, Klumpers DD, Park CY, Rajendran K, Trepat X, van Bezu J, et al. Substrate stiffening promotes endothelial monolayer disruption through enhanced physical forces. Am J Physiol Cell Physiol. 2011 Jan;300(1):C146–54.

